# Identification of soil microbes associated with real-time plastic degradation using *in situ* conductivity sensors

**DOI:** 10.64898/2026.08.16.745074

**Authors:** Andrew J.C. Blakney, Nicole Luna, Nicholas B. Dragone, Taylor Sharpe, Noah Mendez, Kristopher Speetjens, Joshua Garcia, Gregory Whiting, Noah Fierer

**Affiliations:** Cooperative Institute for Research in Environmental Sciences (CIRES), University of Colorado Boulder, Boulder, CO, USA; Mechanical Engineering & Materials Science, University of Colorado Boulder, Boulder, CO, USA; Department of Life & Environmental Sciences, University of California Merced, Merced, CA, USA; Department of Ecology and Evolutionary Biology, University of Colorado Boulder, Boulder, CO, USA

**Author notes:** These authors contributed equally to this work (Andrew J.C. Blakney, Nicole Luna). Corresponding authors: Andrew J.C. Blakney, Noah Fierer.

## Abstract

Microbial-mediated plastic degradation has the potential to address the persistent global problems of plastic waste and pollution. Previous work has shown that soils can harbour microbes capable of plastic degradation, but we expect there is a broader diversity of soil microbes capable of metabolizing plastics than identified to date using more traditional cultivation-based screening methods. Here we demonstrate a novel approach to identify putative plastic degrading microbes in soil. We paired *in situ*, real-time measurements of microbial plastic degradation on conductive sensors with subsequent microbial community profiling of the sensor-associated biofilms exhibiting appreciable degradation. To illustrate the utility of our approach, we focus on microbial degradation of the bioplastic polymer PHBV, poly(3-hydroxybutuyrate-*co*-3-hydroxyvalerate). We screened a range of soils with the *in situ* sensors to identify a subset of five soils with high PHBV degradation rates, and confirmed that PHBV degradation was due to microbial activity. We then extracted DNA directly from sensors placed in soils with high measured rates of PHBV degradation and used marker gene sequencing to identify the bacterial and fungal taxa associated with the observed PHBV degradation. We confirmed via *in vitro* culturing that microbes isolated from the sensors have a demonstrated capacity for PHBV metabolism. Together, these results highlight the benefit and feasibility of using low-cost, in-soil sensors to simultaneously collect real-time data on plastic degradation rates in soil and identify previously unrecognized microbial taxa capable of degrading and metabolizing plastic polymers *in situ*.

## Introduction

“…at present, there is no credible microbial approach to deal with the majority of the world’s accumulated plastic pollution” – Lear *et al*., 2022

Earth has become a plastic planet; with ∼9 billion tonnes of plastic produced globally (Geyer *et al*., 2020), and recycling rates below 10% worldwide (Geyer *et al*., 2017), plastic waste now contaminates terrestrial and aquatic ecosystems across the globe. Plastic contaminants are often resistant to degradation and can have detrimental effects on environmental quality (Brahney *et al*., 2020; Gambarini *et al*., 2021; Galloway *et al*., 2017; Meijer *et al*., 2021), agricultural productivity (Hoffmann *et al*., 2023; Nizzetto *et al*., 2016; Sheng *et al*., 2024), and human health (Landrigan *et al*., 2025; Lear *et al*., 2021), which we are only beginning to document and address (Baho *et al*., 2021; Borrelle *et al*., 2020; MacLeod *et al*., 2021; Simon *et al*., 2021; Vethaak & Legler, 2021; Winton & March, 2025). With global plastic production expected to triple by 2060 (Landrigan *et al*., 2025; Meijer *et al*., 2021; Walker *et al*., 2023), there is a critical need for sustainable technologies capable of degrading plastics at scale to mitigate plastic pollution.

Plastic bioremediation methods based on microbial activities are a promising alternative to current industrial practices for recycling plastics (Kim *et al*., 2022). These microbial-mediated approaches represent a potentially cheaper, lower-energy, and more sustainable alternative to existing industrial waste management strategies (Schade *et al*., 2024). Yet translating this microbial promise into practice remains a major challenge, as no microbially driven process has proven effective at an industrial scale (Gambarini *et al*., 2022). A major barrier to developing microbial-mediated plastic degradation at scale is our limited ability to identify effective plastic degrading microbes in the environment. The vast majority of plastic degrading microbes remain uncultured and functionally uncharacterized (Gambarini *et al*., 2022). To meet our sustainability goals, we need to develop methods that allow researchers to readily identify putative, novel plastic degraders in the environment, and isolate them for the development of innovative bioremediation strategies that fully exploit the plastic degrading potential of microbes.

Soils are a particularly promising target for the discovery of microbial plastic degraders as soil microbial communities are phylogenetically and metabolically diverse (Fierer *et al*., 2017; Gambarini *et al*., 2021), with the capacity to degrade a wide range of recalcitrant organic polymers (Carpena-Istan *et al*., 2025; Gambarini *et al*., 2021). However, the identification of plastic degrading microbes from soils has remained challenging (Viljakainen & Hug, 2021). One reason this challenge persists is because current approaches for identifying plastic degrading microbes are largely untargeted. Most studies screen microbial isolates from arbitrarily selected environmental samples using labour-intensive *in vitro* assays, which are, by definition, restricted to the minority of taxa which are readily culturable, without first determining whether the selected environment exhibits elevated rates of plastic degradation (Danso *et al*., 2019; Dragone *et al*., 2025). Alternatively, we propose that a more effective strategy would be to first identify soils with naturally high plastic degradation rates to reveal previously undescribed microbes which may be effective degraders. We can then use a combination of cultivation-independent and cultivation-dependent approaches to characterize the microbial communities colonizing and degrading the plastic *in situ*. In other words, we want to pair direct, real-time measurements of plastic degradation rates in soil with corresponding analyses of the microbial communities degrading the plastic to identify microbial taxa that hold promise as effective plastic degraders.

To demonstrate the value of this targeted strategy, we focus on the in-soil microbial degradation of a polyhydroxyalkanoate polymer, PHBV (poly(3-hydroxybutuyrate-*co*-3-hydroxyvalerate)). Although PHBV currently only accounts for 1% of global plastic production, it has been touted as a “green polymer” (García-Chumillas *et al*., 2024; Martínez *et al*., 2012), with properties including low toxicity and high UV resistance, strength, and flexibility, that make it a promising alternative to petroleum-based plastics (García-Chumillas *et al*., 2024). Moreover, PHBV is regularly used as a plastic mulch in agricultural systems to enhance crop production (Campanale *et al*., 2024). The first step in our approach is to quantify PHBV degradation rates in soil, which is challenging using more traditional approaches. Current methods, such as mass loss and respirometry, are typically cumbersome, unreliable, and labour intensive, with limited capability for *in situ* testing (Albright & Chai, 2021; Francioni *et al*., 2021). As an alternative strategy, we used electronic sensors placed directly in soil, that can reliably monitor real-time PHBV degradation rates in soil environments, thereby significantly lowering the cost, effort, and time required to identify degradation rates (Atreya *et al*., 2022; Fry *et al*., 2026; Sharpe *et al*., 2026). The sensors have a PHBV trace on one side and a non-biodegradable polymer, PMMA, poly(methyl methacrylate), that serves as an internal control. Conductive graphite particles in the traces give the sensors an initial low resistance value, and as the PHBV is degraded, spacing of the conductive particles increases, leading to a corresponding increase in sensor resistance readings (***Fig. 1 – Step 1***, Atreya *et al*., 2023, Sharpe *et al*., 2026).

**Figure 1.**
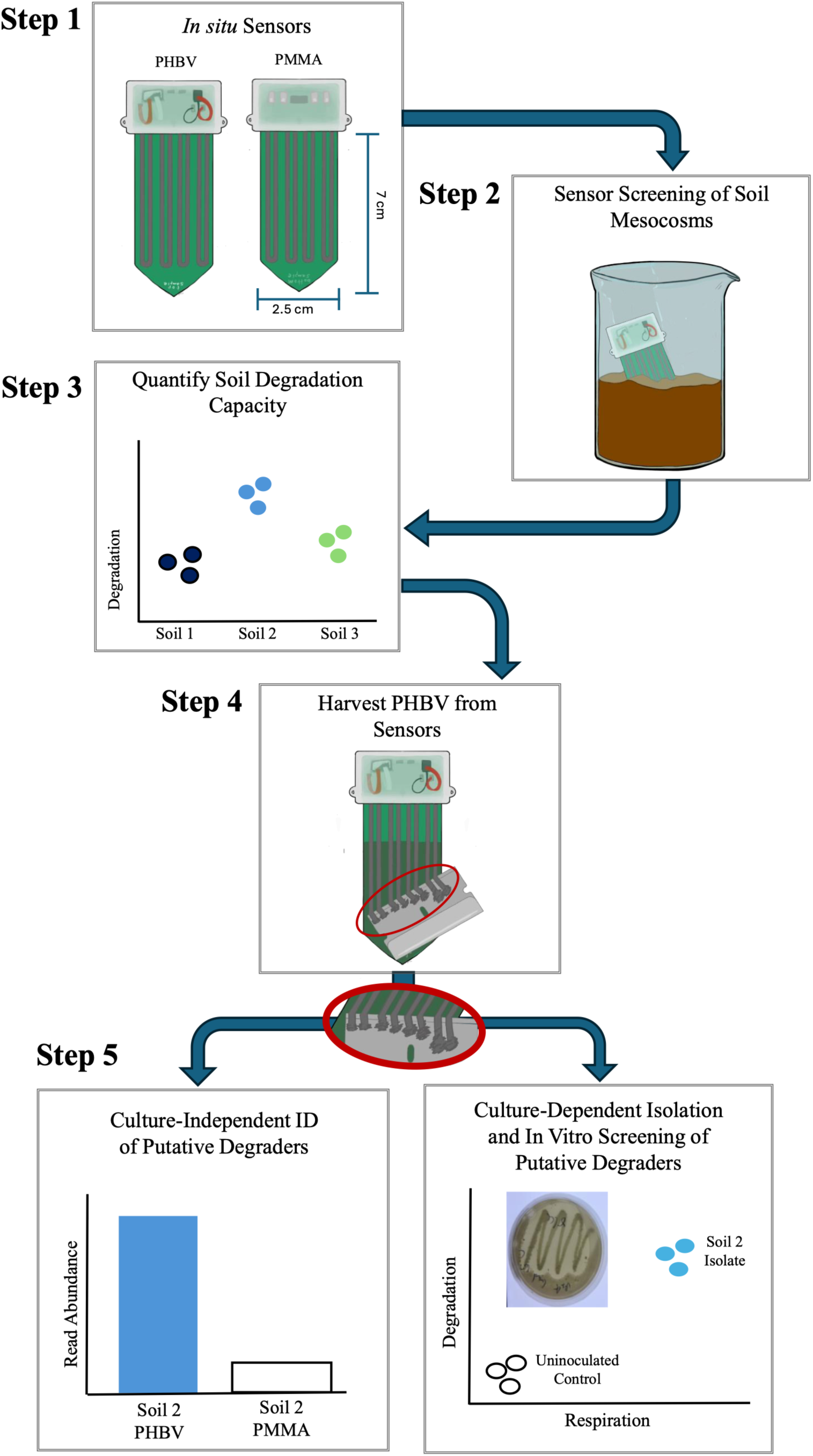
Overview of our strategy to identify novel PHBV degrading microbes from soil using *in situ* electronic sensors. *Step 1*) The sensors are printed with biodegradable PHBV, poly(3-hydroxybutuyrate-*co*-3-hydroxyvalerate), on one side and non-biodegradable PMMA, poly(methyl methacrylate) on the other side as a control. *Step 2*) UV-sterilized sensors are inserted into soil mesocosms and monitor PHBV degradation in real-time as an increase in resistance, which we normalize to the non-degraded PMMA. *Step 3*) At the end of the mesocosm incubation, the sensors are removed and we can summarise the PHBV degradation signal to identify which soils have the highest capacity to degrade PHBV. *Step 4*) PHBV and PMMA material from sensors incubated in the highest PHBV-degrading soils are scraped off and used as starting material for culture-independent (*Step 5*, left panel) and culture-dependent (*Step 6*, right panel) approaches to identify novel PHBV degraders. *Step 5, left*) We used the PHBV and PMMA material from high degrading soils for 16S rRNA and ITS marker gene sequencing to identify microbes enriched on degraded PHBV compared to non-degraded PMMA from each *in situ* sensor. *Step 5, right*) We also used the PHBV material from high degrading soils to isolate novel PHBV degraders on agar plates with a PHBV overlay (inset). Isolates that formed clearing zones on plates were cultured in liquid media to measure their continued capacity to degrade PHBV. We also measured the respiration rates of isolates growing with PHBV as their sole carbon source to confirm PHBV catabolism.

**Figure 2.**
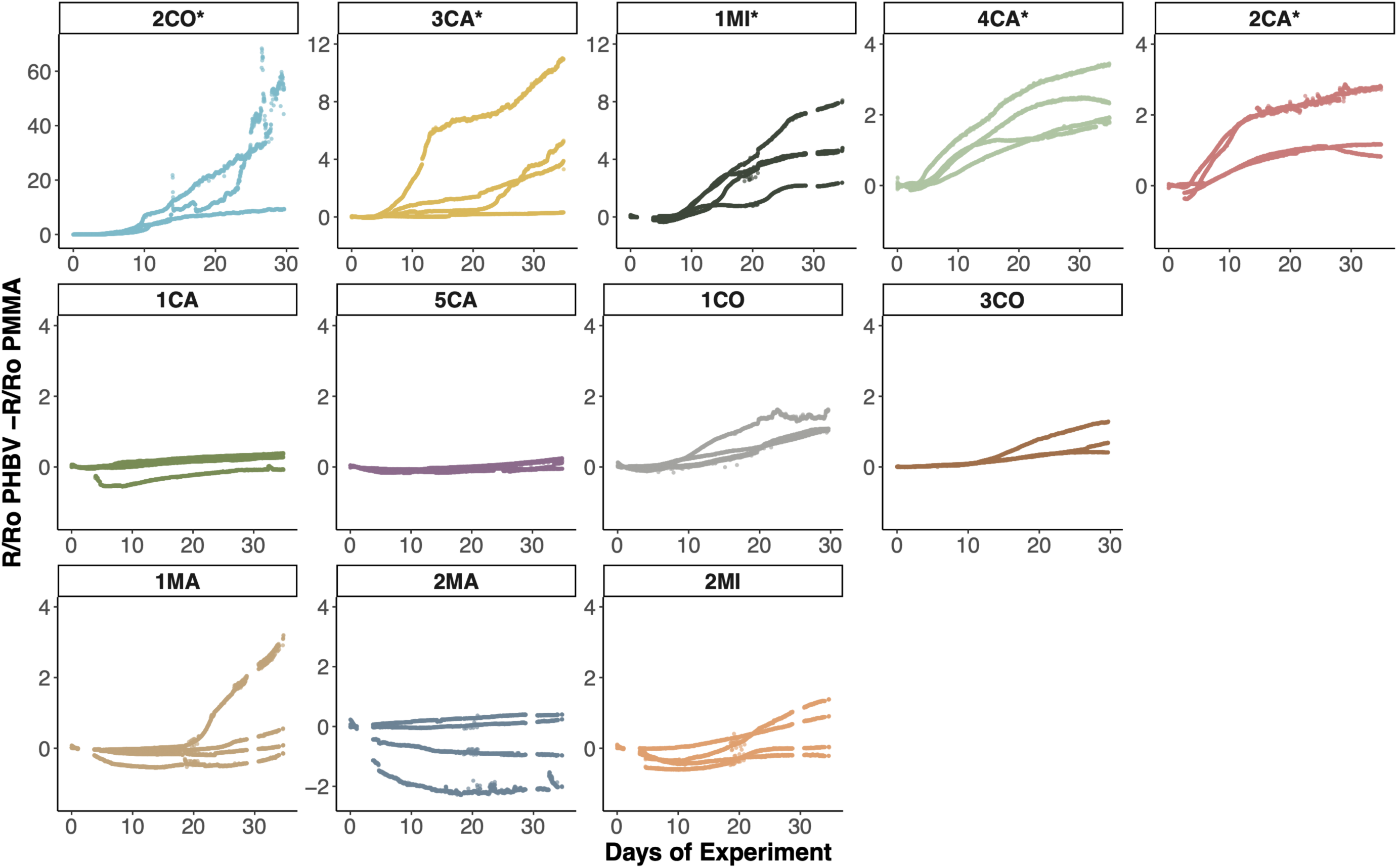
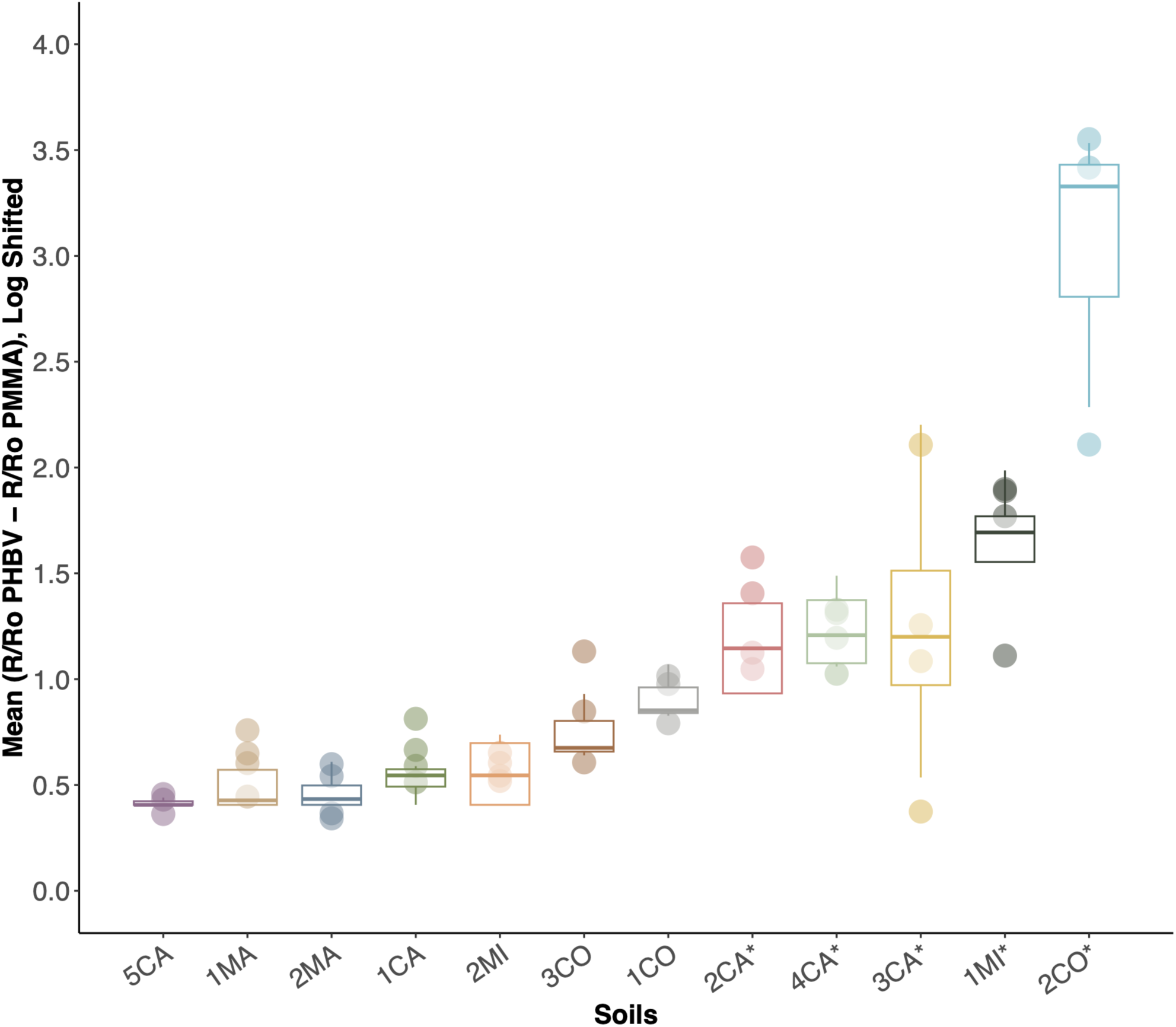
PHBV degradation signals from the 12 screened soils collected from across the USA (CA, California; CO, Colorado; MA, Massachusetts; MI, Michigan). We selected the five soils with the highest PHBV degradation capacity (2CA, 4CA, 3CA, 1MI, and 2CO) for downstream microbial analyses. A) Change in PHBV degradation signals over time across the 5-week incubation period. B) Day 19.5-29.5 summarized PHBV degradation sensor signal for soil mesocosms, as the highest signal was recorded from this timeframe due to accumulated degradation. N = 4 for all soil mesocosms presented here, except 1CO, 2CO, 3CO, and 5CA, which had 3. We removed one 5CA sensor that had mechanical damage.

Here, we first screened 12 distinct soils using these PHBV-PMMA sensors, with paired sterilized soils to confirm that the measured PHBV degradation was a product of microbial activities (***Fig. 1 – Step 2***). From this screening effort, we identified a subset of five soils where degradation rates were highest (***Fig. 1 – Step 3***), and selected these soils for subsequent microbial analyses. We then used a cultivation-independent approach, isolating DNA directly from both the degraded PHBV and non-degraded PMMA on each *in situ* sensor (***Fig. 1 – Step 4***). We analyzed these DNA samples via marker gene sequencing to identify bacterial and fungal taxa enriched on degraded PHBV compared to PMMA (***Fig. 1 – Step 5***, left panel). Given that not all PHBV-associated taxa are necessarily capable of degrading PHBV, we complemented the culture-independent work with cultivation-dependent analyses to confirm that a subset of taxa can degrade PHBV *in vitro* (***Fig. 1 – Step 5***, right panel). While this work focused on a single plastic type in a relatively small subset of soils, this approach could ultimately be expanded to include other plastic polymers, and environmental conditions, to directly measure and characterise microbial-mediated plastic degradation *in situ* and accelerate the search for effective plastic degrading microbes.

## Materials and Methods

### Sensor Construction

To identify *in situ* soil PHBV degradation, we used electronic sensors similar to those which have been described previously (Atreya *et al*., 2023; Fry *et al*., 2026; Sharpe *et al*., 2026). Briefly, the active PHBV ink was prepared as described by Atreya *et al*. (2023), and composed of 1 g of biodegradable PHBV (Sigma-Aldrich, cat. no. 403105), with 8% mol hydroxyvalerate content, 12 mL of chloroform (Sigma-Aldrich), and 3 g of 7-11 µm graphite powder (Thermo Scientific, cat. no. 046304.30). The control PMMA ink was prepared in the same fashion, with 0.94 g of PMMA (Sigma-Aldrich, cat. no. 445746), 12 mL of chloroform, then 3 g of 7-11 µm graphite. The sensor substrate was a printed circuit board designed for 4 “U” shaped PHBV traces connected in a parallel circuit on the front, and 4 identically shaped PMMA traces connected in a separate parallel circuit on the back (***Fig. 1 – Step 1***). The ink was applied to the circuit board via a stencil printing process, as described by Atreya *et al*. (2023), with the PMMA traces printed and dried, followed by the PHBV traces. Two cables were soldered onto the circuit board for PHBV and PMMA trace resistances to be measured independently, and the contacts weatherized (Fry *et al*., 2026), leaving a final 3.8 mg of PHBV trace exposed for degradation. We confirmed that UV sterilization (10 minutes per side) did not impact sensor readings, with the sterility of both sides of the sensors confirmed before installation.

### Soil Collection and Preparation

To identify soils with a high capacity for PHBV degradation, we screened 12 different soils (0 – 10 cm depth) collected from across the United States. These soils were selected to span a range of site and edaphic characteristics (***Supp. Table 1***). Soils were transported on ice in coolers back to the lab, where they were sieved to 2 mm, homogenized and then stored at

4°C prior to use. Subsamples were taken from each soil for chemical analyses (Ward Labs, Kearny, NE), and triplicate subsamples of each bulk soil were collected for DNA extraction, amplicon sequencing and microbial community analyses, as described below. We determined the existing water content and water holding capacity (WHC) of each soil to standardize each soil mesocosm to 50% WHC.

We measured soil respiration rates, as a proxy for microbial activity, using the alkaline trap method (Jensen *et al*., 1996; Yim *et al*., 2002). Briefly, triplicate 5 g subsamples of each soil were adjusted to 50% WHC for a 1-week equilibration period, after which an alkaline trap of 1 M NaOH in a 1.5 mL tube was placed in each soil sample, and left to incubate for 2 weeks (only 1 week for soils 1CA and 5CA, as they saturated the trap with longer incubation) in the dark at room temperature. Triplicate, soil-free blank tubes were also included to measure ambient CO_2_ concentrations.

### Mesocosm Experimental Setup and Sampling

To screen the 12 soils for PHBV degrading capacity, and confirm that degradation was microbially mediated, we prepared live and sterile soil mesocosms with subsamples from each of the 12 soils (***Fig. 1 – Step 2***). Under aseptic conditions, we weighed 300 g of triple autoclaved subsamples of each soil into sterile mesocosms (King *et al*., 2024), followed by 300 g of live soil into their respective mesocosms. We then adjusted the gravimetric WHC of each mesocosm to 50% using sterile dH_2_0.

We inserted one UV-sterilized PHBV-PMMA sensor into each mesocosm (***Fig. 1 – Step 2***), including 8 sterilized blank mesocosms without any soil, for a total of 80 mesocosms (12 soil types *6 replicates +8 blanks). Each mesocosm was sealed with a double layer of autoclaved aluminium foil. We recorded the final weight of each mesocosm to monitor potential water loss. We randomized the mesocosms in a large plastic bin, lined with damp paper towel to minimize soil drying. The mesocosms were incubated at room temperature (approximately 23°C) for up to 5 weeks.

We terminated the experiment within 5 weeks to balance true degradation in the live soil mesocosms, versus signal caused by potential grow back in the sterilized soil mesocosms. Sensor resistance readings were acquired via custom 8-channel dataloggers at 30-minute intervals. Dataloggers were constructed with a WiFi-enabled microcontroller (Adafruit ESP32 Feather V2) and powered with lithium polymer batteries through a low-power timer circuit (Sharpe *et al*., 2026). We harvested the mesocosms by first disconnecting each sensor from its datalogger, then under aseptic conditions, removing each sensor from the sterile mesocosms, and placing them individually in sterile WhirlPak bags (Nasco Sampling, WI). We then repeated sensor harvesting from the live mesocosms. Sensors were stored at –20°C until further processed.

### Sensor Resistance Data Analysis

The sensor trace resistances measured by the dataloggers indicated the real-time degree of degradation of PHBV in each mesocosm. These resistance readings can be predictably and reversibly affected by soil conductivity, as the soil provides an alternate current path (Sharpe *et al*., 2026). We normalized each trace resistance to its average resistance in the 0.5-1.5 day period after installation in the mesocosms to account for initial differences in soil conductivity, (Fry *et al*., 2026; Sharpe *et al*., 2026). Any background signal drift associated with potential changes in soil conductivity, or other parameters, that might occur over the course of the incubation period can be accounted for with the PMMA traces, which should not be degraded by microbial activity in soil (Thakore *et al*., 2001). PMMA traces have previously shown no signs of degradation via change in resistance or microscopy images after incubation in compost tea (Atreya *et al*., 2023). Since changes in resistance readings of PMMA traces are primarily due to changing soil conductivity and the PMMA and PHBV traces have matched geometry and initial resistivity, a change in soil conductivity will create a similar change in resistance reading to both traces. Thus, the non-biodegradable PMMA trace signal can be used to subtract out any abiotic signal drift by calculating the normalized PHBV trace resistance minus the background signal of normalized PMMA trace resistance (R/R_o_ PHBV – R/R_o_ PMMA). To compare cumulative degradation rates across the soils, we calculated the mean (R/R_o_ PHBV – R/R_o_ PMMA) over the last ten-day window of the incubation period (days 19.5-29.5).

### DNA Extraction and 16S rRNA and ITS Marker Gene Sequencing

We selected five soil types that exhibited the highest cumulative rates of PHBV degradation to characterize the microbial communities on the sensors in the unsterilized mesocosms and in the surrounding bulk soil (***Fig. 1 – Step 3***). Excess soil was gently removed from the sensors while still frozen. Under aseptic conditions, PMMA was first scraped off the sensor with a sterile razor and collected in a sterile 1.5 mL microcentrifuge tube (***Fig. 1 – Step 4***). The scraping process was then repeated for the residual PHBV trace on the other side of the same sensor. This process yielded two trace samples per sensor for a total of 46 samples (n = 23 per substrate, PHBV and PMMA). All samples were stored at –80°C prior to DNA extraction.

Total genomic DNA was extracted directly from each sample (36 bulk soils +46 trace substrates) using DNeasy Plant DNA Extraction Kits (Qiagen, Germany) as previously described (Blakney *et al*., 2024). We included 7 DNA extraction blanks as negative controls to test for potential contamination. To characterize the prokaryotic and fungal communities on the PHBV and PMMA trace material and the bulk soil samples, extracted DNA from all samples were used to prepare amplicon libraries as previously described (Blakney *et al*., 2024 & 2025). First, 25 μL of DNA from each sample were transferred to 96-well plates with well locations randomized. We included 3 no-template negative controls to check for potential contamination introduced during downstream processing. DNA samples were used as templates for PCR amplification for both the 16S rRNA gene and ITS regions, using the primers (515f-806r and ITS1-F-ITS2-R), with the PCR recipes and amplification protocols described in Winfrey *et al*. (2025). We included 2 no-template PCR blanks in each reaction to check for the introduction of potential contaminants at the PCR step. All PCR reactions were run in duplicate, and successful amplification confirmed via gel electrophoresis. Amplicons from the duplicate reactions were then combined prior to clean-up and normalization, following a protocol described previously (Buchner, 2022). Finally, we sequenced both marker gene libraries separately on an Oxford Nanopore Technologies (ONT) MinION, with libraries prepared using the ONT Ligation Sequencing Kit V14.

### Bioinformatics Analyses

The integrity of the sequence data was confirmed using MD5 checksum protocol (Roy *et al*., 2018). Subsequently, all data was managed and analysed in R (4.4.2 R Core Team, 2024), and plotted using ggplot2 (Wickham, 2016). We obtained 22 305 734 and 42 825 865 sequencing reads from the 16S rRNA and ITS amplicon pools, respectively, which are publicly available at NCBI Bioproject PRJNA1456121, along with all corresponding metadata.

Processing was completed as previously described (Gebert *et al*., *In Press*) to remove low quality reads (> Q = 20) with chopper (De Coster & Rademakers, 2023), and using cutadapt (Martin, 2011) to remove reads < 200 bp in size. We retained 3 061 035 and 6 052 550 reads for 16S rRNA and ITS ASV (amplicon sequence variants) inference with DADA2 (Callahan *et al*., 2016), which generated 20 749 prokaryotic ASVs and 6741 fungal ASVs across the whole dataset, including both bulk soils and sensor trace samples. We used the default settings throughout the DADA2 pipeline, except the dada inference function, which used the pool =‘pseudo’ argument to increase the likelihood of identifying rare taxa. Consequently, the chimera removal function also included the method = ‘pooled’ argument, as previously described (Blakney *et al*., 2024). We assigned taxonomy against the SILVA database for 16S rRNA ASVs (nr99_v138.1; Yilmaz *et al*., 2014), and the UNITE database for the ITS ASVs (19.02.2025; Abarenkov *et al*., 2022). ASVs, taxonomy, and metadata were organized into prokaryotic and fungal phyloseq objects for downstream analysis (McMurdie & Holmes, 2013).

We then filtered the phyloseq objects to remove prokaryotic ASVs identified as chloroplast and mitochondria, or any samples that had fewer than 250 reads in total. This processing removed all the control samples, and one experimental sample (1MI). We then removed any ASV that had fewer than 10 reads across the entire dataset, which included any ASVs identified in the controls. This process left all samples with more than 2000 reads, which were assigned to 18 157 prokaryotic ASVs. We filtered the fungal data by removing any samples with fewer than 1000 reads, which again removed the controls, and the same experimental sample (1MI), followed by removing any ASVs that had fewer than 10 reads in total. This left 5398 fungal ASVs, and all samples with more than 1800 reads.

To test if the microbial communities differed between the bulk soils, we used a non-parametric permutational multivariate ANOVA (PermANOVA). After Hellinger transformation (Legendre & De Cáceres, 2013), a distance matrix was calculated using Bray-Curtis dissimilarity. We visualized microbial community differences among the soils using a principal coordinate analysis, and relative abundance plot. We then tested if the microbial communities differed between sample types (i.e. bulk soil, PHBV trace, and PMMA trace) within each of the five PHBV-degrading soils. Data were rarefied by soil type, then tested by PermANOVA, and plotted as described above.

We then identified which microbes were enriched on PHBV compared to PMMA (***Fig. 1 – Step 5***, left panel) in the high PHBV degrading soils. First, bulk soil samples were removed from each phyloseq object, which were then separated and rarefied by each soil type. Next, within the dataset for each soil type, we filtered our PHBV read abundance data such that each prokaryotic or fungal ASV retained needed to represent at least 0.1% of reads in each PHBV sample, and at least 0.05% of total PHBV reads. This left 3-24 bacterial and fungal ASVs per soil type that were highly abundant on PHBV. We then calculated log2-fold change in abundances between the PHBV and PMMA reads for each of these PHBV-abundant ASVs, adding a pseudocount of 1 to account for read abundances of 0.

### Culture-Dependent Identification of PHBV Degrading Microbes

We used a screening method described in Dragone *et al* (2025) to test the PHBV degradation capacity of culturable microbes from the PHBV traces (***Fig. 1 – Step 5***, right panel). Briefly, putative PHBV degrading microbes cultured from PHBV trace material were identified through the production of a clearing zone in PHBV overlay on R2A plates. We serially diluted ∼1 mg of the same PHBV trace used for DNA extraction from the 2CO soil in 1X PBS, where 50 µL of each dilution (10^-1^ – 10^-6^) was spread onto the PHBV-R2A plates in triplicate. Three “blank” PHBV-R2A plates were inoculated with 50 µL of 1X PBS to account for potential contamination introduced during media prep and plating. All plates were incubated at 25°C for four weeks under aerobic conditions and were checked weekly for clearing zones in the PHBV overlay (see ***Fig. 5*** for example). Colonies that created clearing zones were re-streaked at least three times on new PHBV-R2A plates to generate axenic cultures and confirm PHBV degradation ability.

We identified the PHBV degrading isolates via Sanger sequencing of the full-length, ∼1400 bp archaeal and bacterial 16S rRNA gene (27F and 1492R primer pair, Frank *et al*., 2008; Weisburg *et al*., 1991), as described in Dragone *et al* (2023). DNA was extracted from each isolate using Qiagen’s DNeasy Ultraclean Microbial Kit (Qiagen, Germantown, MD, US) following the manufacturer’s instructions. DNA from each isolate was used as template for PCR amplification, following the PCR recipe and protocol as described above. Amplified PCR product from all the cultured isolates, six DNA-extraction blanks, and three no-template PCR blanks were sequenced by Azenta Life Sciences’ Genewiz Sanger sequencing service (Azenta Life Sciences, Burlington, MA, USA).

Whole genome sequencing was performed as described in Dragone *et al*., (2026). DNA aliquots from two of our isolates, *Streptomyces sp.* [2CO Garden Isolate], *Lysinibacillus sp.* [2CO Garden Isolate] were sequenced by Plasmidsaurus using their bacterial genome sequencing service. Raw sequencing data was then filtered with Filtlong v.0.2.1 (https://github.com/rrwick/Filtlong) following default parameters to remove the 5% lowest quality reads. Filtered reads were then assembled with Flye v.2.9.5 (Kolmogorov *et al*., 2019) with parameters for high quality ONT reads. The generated assemblies were polished with Medaka v.2.0.1 (https://github.com/nanoporetech/medaka) and Polypolish v.0.6.0 (Bouras *et al*., 2024; Wick & Holt, 2022). Assembly quality was assessed with CheckM2 (Chklovski *et al*., 2023), open reading frames were predicted with Prodigal v.2.6.3 (Hyatt *et al*., 2010), and taxonomy was assigned with the GTDBtk v.2.1.0 classify workflow (Chaumeil *et al*., 2020) based on the Genome Taxonomy Database v.226 (Park et al., 2022). Identification of plastic degradation genes in each genome was performed by comparing the translated amino acid sequences against the PlasticDB database (Gambarini *et al*., 2022).

### Testing Sensor PHBV Degradation Corresponds to Microbial Catabolism of PHBV

To further confirm that the PHBV degradation signal from the *in situ* sensors reflected microbial PHBV degradation and catabolism of PHBV to CO_2_ (***Fig. 1 – Step 5***, right panel), we set up controlled experiments with new PHBV-PMMA sensors and CO_2_ respirometers (OXMAN, New York, NY, USA). Each sensor was placed at the bottom of a different 500 mL gooseneck Erlenmeyer flask (six total) containing 250 mL of sterile M9 media, with the wires threaded out of the neck opening, which was then sealed to make the neck opening gastight.

We grew three 50 mL cultures of isolate *Lysinibacillus sp.* [2CO Garden Isolate] overnight in R2A broth. Cultures were grown up to an OD₆₀₀ of 1.04 (biomass equivalent). These overnight cultures were centrifuged at 10,000g for 30 minutes to concentrate cells, and all R2A was pipetted out. The concentrated cells were washed three times with 10 mL of 1X PBS buffer to remove excess R2A media, and then each tube of washed cells was then resuspended in 5 mL of M9 media. We then removed 5 mL of M9 media from three of the Erlenmeyer flasks, which were then inoculated with the resuspended cells (‘Live’ treatment). The remaining three flasks were left uninoculated as sterile controls. Flask-mounted CO₂ respirometers were attached to each flask with teflon tape to ensure an airtight seal. The respirometers were calibrated to an atmospheric CO_2_ concentration of 430 ppm, and set to record headspace CO_2_ concentration every 10 seconds, and pump the headspace of each flask so that CO_2_ concentrations never exceeded 2000 ppm (pump would bring CO_2_ concentrations to 1000 ppm).

Flasks were left to incubate at room temperature on a shaking plate for 30 days. After this incubation, sensor readings and CO_2_ production readings were downloaded. Sensor readings were processed following methods described above. CO_2_ production above the atmospheric baseline was calculated by integrating the raw respirometer reading per hour using the R package ‘pracma’ (Borchers, 2025). Material scraped from the live sensors was re-identified with Sanger sequencing of the 16S rRNA gene (see above) to confirm the identity of the isolate upon completion of the experiment.

## Results and Discussion

### PHBV degradation rates varied across soil types

To identify how soils vary in their PHBV-degrading capacity, we screened 12 soils collected from across the continental United States using the *in situ* sensors (***Fig. 1 – Step 2***), with these soils representing a broad range in site and edaphic characteristics (***Supp. Table 1***). For example, pH values ranged from 5.1 to 7.4, percent organic carbon ranged from 1.2 to 14.1, the rates of microbial activity (total CO_2_ production over 1 week) ranged from 4.05 to 26.62 µg C-CO2 g soil^-1^ day^-1^. Unsurprisingly, the overall composition of the bulk soil prokaryotic (i.e. bacterial and archaeal) and fungal communities also varied across the 12 soils (***Supp. Fig. 1B***), which was also evident by comparing the relative abundances of major taxonomic groups ***(Supp. Fig. 1A)***.

Across the 12 soils, we also observed appreciable variation in measured PHBV degradation rates (***Fig. 2A***). Mean PHBV degradation per soil over the last ten days of the incubation period—i.e., the timeframe of highest signal—varied from 0.018 to 22.2, with the 2CO soil having the greatest capacity for PHBV degradation (***Fig. 2B***). In seven of the 12 soils screened, measured PHBV degradation rates were negligible, or inconsistent across replicate mesocosms; the remaining five soils with the highest PHBV degradation were selected for downstream microbial analyses (***Fig. 2B***). Our observation that the capacity for PHBV degradation is so variable across soils (***Fig. 2B***) is consistent with previous studies even when soils are incubated at the same temperature and moisture levels (Meng *et al*., 2023). The measured differences in PHBV degradation across the 12 soils (***Fig. 2B***) were not correlated with variation in any of the measured soil edaphic properties (***Supp. Table 1; Supp. Fig. 2***). This was true even for observed prokaryotic, and fungal species richness, and measured rates of total microbial activity (microbial CO_2_ production), which were not correlated with measured rates of PHBV degradation (***Supp. Fig. 2***). The soils with the highest PHBV degradation capacity were not necessarily those soils with the highest rates of microbial activity and PHBV degradation capacity cannot be effectively predicted from edaphic variables alone. Instead, we expect that it is the amounts and types of active PHBV degrading microorganisms in soil, not abiotic characteristics, or overall levels of microbial activity, that largely determine the capacity for PHBV degradation in any given soil.

Although previous experiments with these sensors have demonstrated that the sensor signal reflects biodegradation of PHBV, not degradation by abiotic processes (Atreya *et al*., 2023), we wanted to confirm that the results presented in ***Fig. 2*** reflect degradation of PHBV via microbial processes. There are three lines of evidence to suggest that this is indeed the case. First, we included mesocosms of sterilized soils for each soil type. For soils that had PHBV degradation signals from ‘live’ mesocosms, the PHBV degradation from sensors in the corresponding sterilized mesocosms were typically far lower (***Supp. Fig. 3***). For the subset of sensors placed in sterilized soils where a signal response was observed, the signal was only evident towards the end of the incubation period (***Supp. Fig. 3A***), likely due to microbial growth after sterilization, a common issue even in triple autoclaved soils (King *et al*., 2024). Second, further evidence that the sensors were detecting microbial degradation, and not degradation by abiotic processes, comes from the *in vitro* assays with plastic degrading isolates, as discussed in more detail below. Finally, we included non-degradable PMMA traces on each sensor to account for background soil abiotic factors, particularly conductivity (Atreya *et al*., 2023; Thakore *et al*., 2001). Therefore, given that we observed no abiotic degradation from the PMMA traces, and minimal activity of the sensors in sterile soils, the majority of PHBV degradation signal observed in the live soils presumably reflects PHBV degradation via microbial activity.

**Figure 3.**
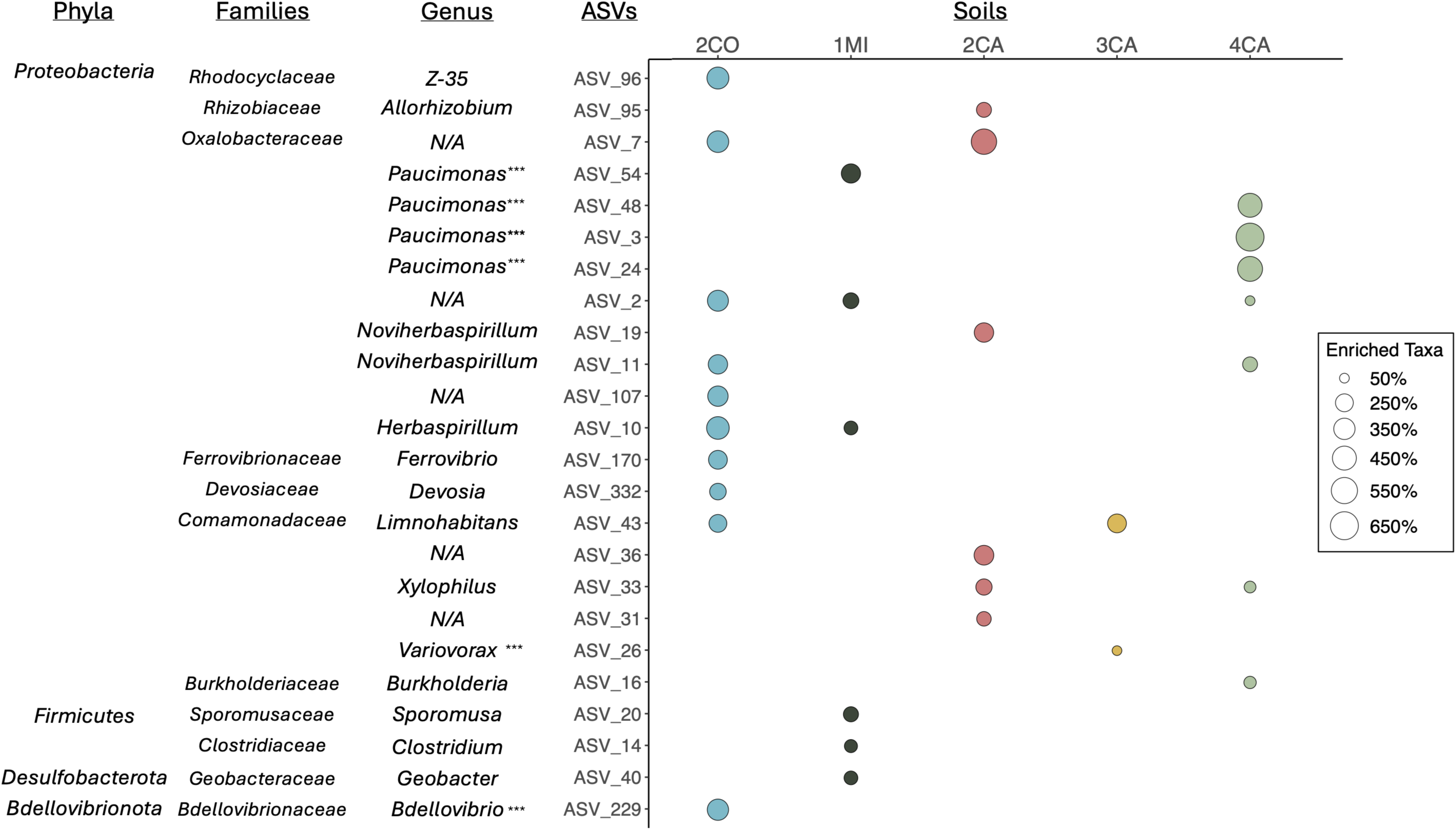
Bacterial ASVs enriched on PHBV were identified from the PHBV trace materials of five, high PHBV degrading soils. Enriched ASVs represent those which were 50% or more abundant on PHBV compared to the non-degradable PMMA control (i.e. 2^Log2FC^ > 2^0.585^) with the legend indicating the magnitude of enrichment. ASVs marked with asterisks represent taxa that have previously been identified as being capable of plastic degradation (Gambarini *et al*., 2022).

### Identification of bacteria and fungi enriched on PHBV

We next compared the microbial communities associated with the bulk soils versus the PHBV and PMMA traces by marker gene sequencing of bacterial and fungal communities, focusing on the subset of five soils with the highest observed PHBV degradation rates (***Fig. 2B***). Note that the DNA samples used for these cultivation-independent analyses were extracted directly from the sensor trace material or the bulk soils (***Fig. 1 – Step 4***). We found that, for each of the five soils, the overall composition of the bacterial and fungal communities found on PHBV or PMMA traces was distinct from the corresponding bulk soil communities (***Supp. Fig. 4***). These results indicate that the sensor surfaces enrich for unique subsets of the bulk soil communities.

**Figure 4.**
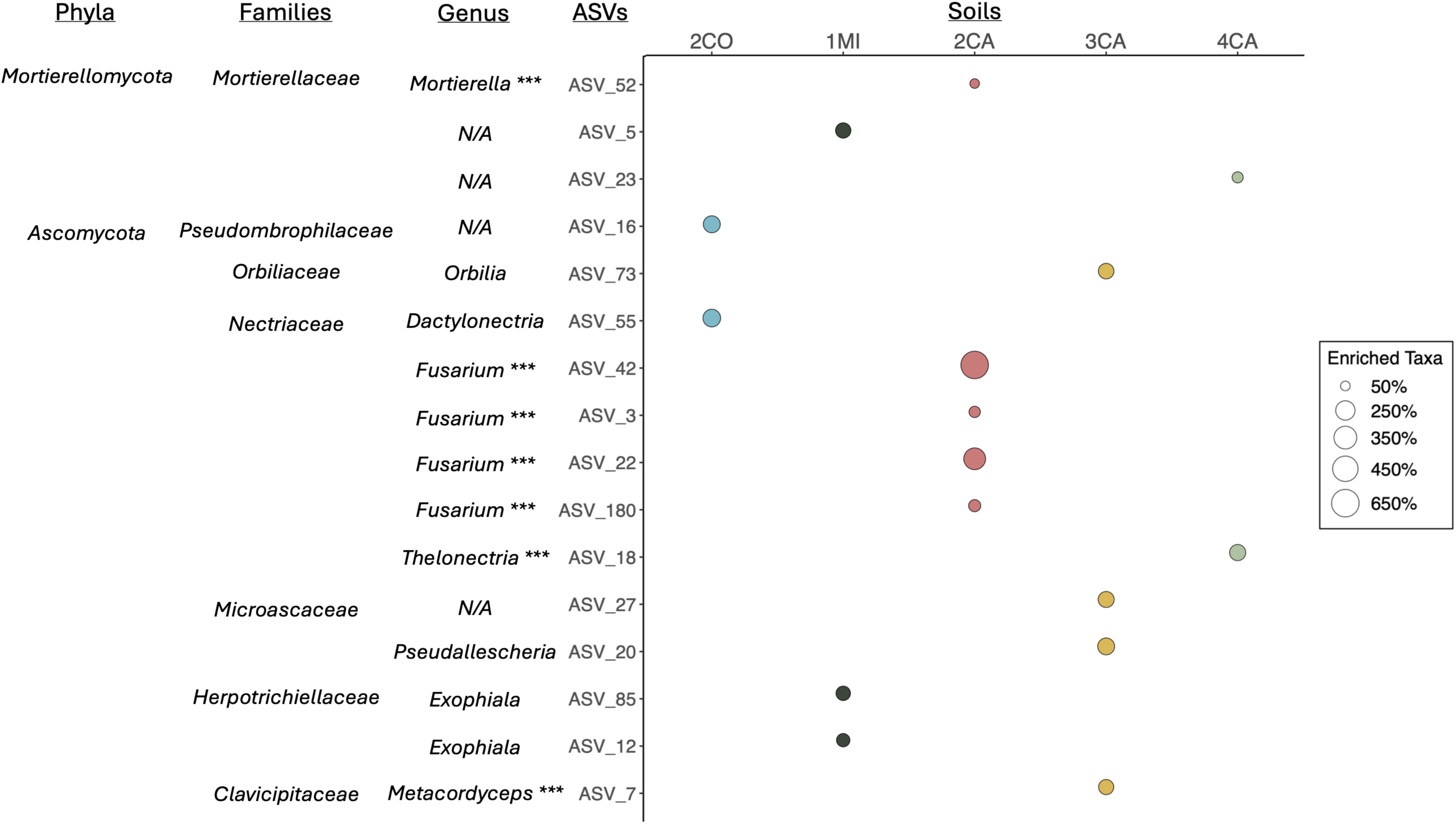
Fungal ASVs enriched on PHBV were identified from the PHBV trace materials of five, high PHBV degrading soils. Enriched ASVs represent those which were 50% or more abundant on PHBV compared to the non-degradable PMMA control (i.e. 2^Log2FC^ > 2^0.585^) with the legend indicating the magnitude of enrichment. ASVs marked with asterisks represent taxa that have previously been identified as being capable of plastic degradation (Gambarini *et al*., 2022).

**Figure 5.**
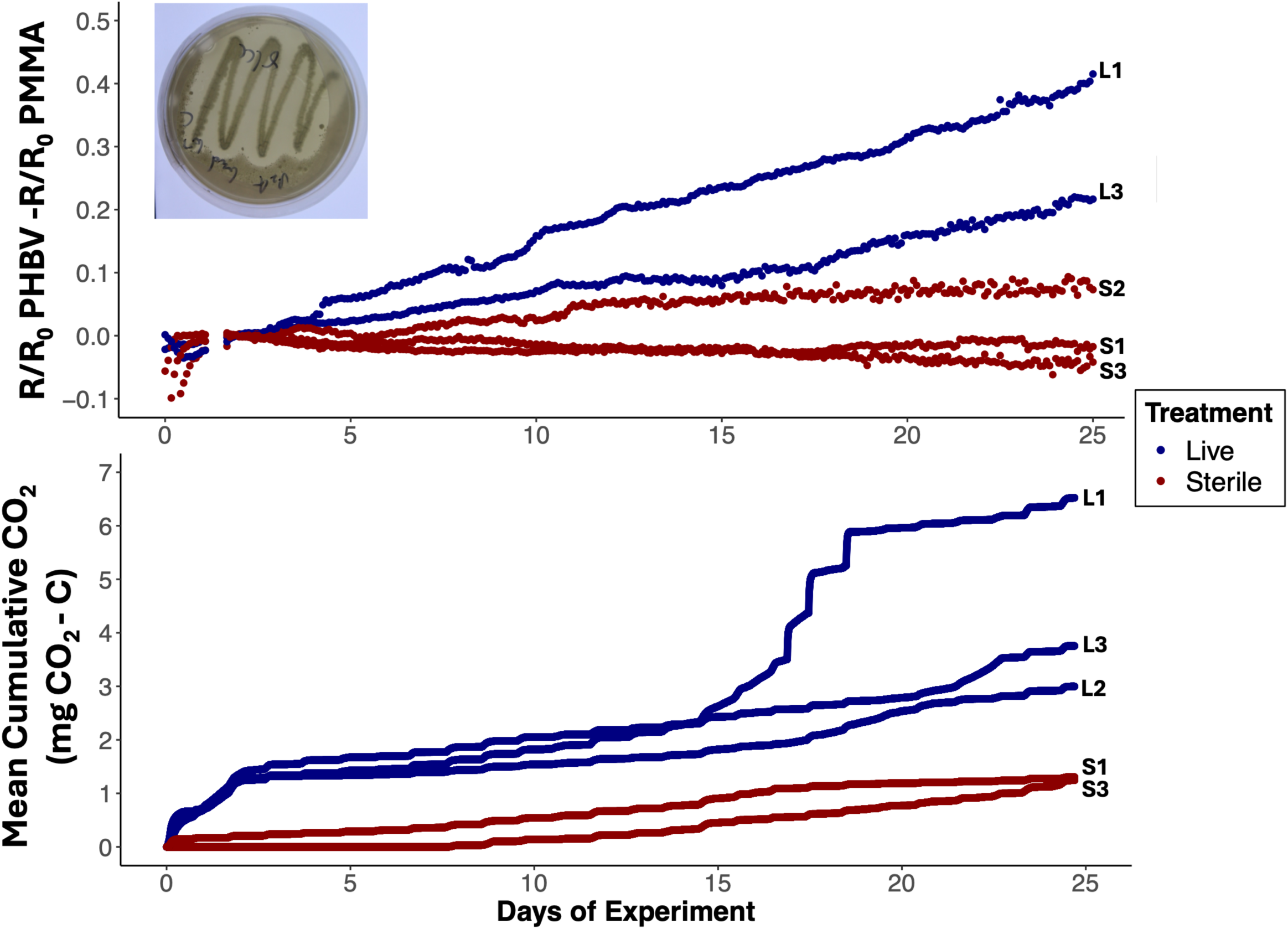
*Lysinibacillus* sp. isolated from the high PHBV degrading 2CO soil on R2A plates with a PHBV overlay (inset) maintains its capacity to degrade PHBV in M9 culture where the PHBV-PMMA sensor is the only carbon source (top panel). Flask-mounted respirometers recorded cumulative CO_2_ production as concurrent with PHBV degradation (bottom panel). Uninoculated controls had no PHBV degradation nor CO₂ production throughout the experiment. The L2 PHBV-PMMA sensor was removed due to damage (top panel), and the CO_2_ data from S2 was removed due to respirometer malfunction (bottom panel).

We next sought to identify the specific prokaryotic or fungal taxa enriched on the degraded PHBV as compared to the corresponding non-degradable PMMA trace to identify taxa associated with the observed PHBV degradation (***Fig. 1 – Step 5***, left panel). This comparison of the PHBV versus the PMMA communities within a given soil type allows us to distinguish between those taxa that are merely adhering to the sensor surfaces from those that are more likely to be responsible for PHBV degradation. We were able to identify a total of 24 bacterial ASVs that were enriched on PHBV across the five soils (***Fig. 3***). Some of these ASVs were only identified from a single soil, but a subset were identified across multiple soils, including ASVs within the families *Comamonadaceae*, such as *Limnohabitans* and *Xylophilus* spp., and the *Oxalobacteraceae*, including members of the genera *Paucimonas*, *Noviherbaspirillum*, and *Herbaspirillum*. The enriched putative PHBV-degraders spanned four phyla, 11 families, and most (18/24) have not been previously identified to degrade any type of plastic (Gambarini *et al*., 2022), highlighting the potentially untapped metabolic potential of soil bacteria to degrade PHBV. *Oxalobacteraceae* and *Comamonadaceae* (both *Proteobacteria*) were highly represented as PHBV-enriched ASVs. Taxa within the genera *Paucimonas* (*Oxalobacteraceae*) and *Variovorax* (*Comamonadaceae*) have previously been shown to degrade PHBV (Gambarini *et al*., 2022; Martínez-Tobón *et al*., 2018; Mergaert *et al*., 2000; Suyama *et al*., 1998). Isolates in the genus *Variovorax* have also been shown to degrade a variety of other plastics (Gambarini *et al*., 2022; Mergaert *et al*., 2000; Suyama *et al*., 1998), while species identified as *Bdellovibrio* have been shown to degrade polyhydroxyalkanoates (Gambarini *et al*., 2022; Martínez *et al*., 2012).

Across all five soils, we identified a total of 16 fungal ASVs as putative PHBV-degraders, spanning two phyla, and seven families. None of these enriched fungal ASVs were shared across the five soils, and half the taxa have not been previously identified to degrade any type of plastic (Gambarini *et al*., 2022). Three genera, *Thelonectria*, *Metacordyceps*, and *Fusarium*, have isolates shown to degrade polyurethanes (Gambarini *et al*., 2022; Mergaert *et al*., 1992; Navarro *et al*., 2021). Taxa within the *Fusarium* genus have also been shown to degrade a wide variety of other plastics, including PHB (Abe *et al*., 2010; Arefian *et al*., 2013; De Hoe *et al*., 2018; Gambarini *et al*., 2022; Jeszeová *et al*., 2018; Murphy *et al*., 1996; Navarro *et al*., 2021; Sang *et al*., 2002; Zumstein *et al*., 2017). Isolates from the *Mortierella* and *Dactylonectria* genera have been demonstrated to degrade other types of plastic polymers (Gambarini *et al*., 2022; Jeszeová *et al*., 2018; Koutny *et al*., 2006).

Together these results demonstrate that we were able to identify both bacterial and fungal taxa that may be associated with PHBV degradation as they were enriched on the PHBV sensor traces. These taxa included those previously demonstrated to be capable of PHBV degradation as well as novel taxa which have not previously been screened for their plastic degrading potential. Importantly, we show that while some bacterial taxa are shared across soils, many of the taxa found to be enriched on PHBV are unique to individual soils, highlighting that plastic degrading microbes can be soil-specific and the microbial taxa that colonized PHBV differ depending on the soil in question, results that parallel those observed previously (Bandopadhyay *et al*., 2020; Liu *et al*., 2026; Rauscher *et al*., 2023; Rüthi *et al*., 2020; Yu *et al*., 2023). We also identified a broad diversity of novel fungal taxa associated with PHBV degradation. The capacity for plastic degradation by fungi is likely higher than previously known given that fungi are generally under-represented in efforts to screen for plastic degrading microbes using cultivation-dependent approaches (Gambarini *et al*., 2021). Our results also suggest that plastic degradation in environmental samples is unlikely to be a product of individual taxa degrading PHBV on their own but is more likely a result of diverse microbial taxa colonizing plastic substrates (Datta *et al*., 2016; Dey *et al*., 2022; Wright *et al*., 2021).

These cultivation-independent analyses come with some important caveats. The relative importance of the individual bacterial and fungal taxa to PHBV degradation remains unknown. Likewise, we cannot confirm from these data alone that those taxa which were identified as being enriched on PHBV versus the corresponding PMMA trace necessarily have the capacity for PHBV degradation. Rather, some taxa may simply be associated with other taxa that are directly responsible for PHBV degradation, or metabolizing other substrates, and just adhering to the PHBV trace. However, given that some taxa known to be capable of PHBV degradation were identified with our cultivation-independent analyses, our strategy of pairing measurements of *in situ* PHBV degradation with analyses of the communities on the degraded sensors demonstrates the utility of this approach for the exploration of novel plastic-degrading microbes in soil environments.

### Characterization of bacterial isolates cultivated from the sensors

To further demonstrate the utility of the *in situ* PHBV sensors, and complement our culture-independent analyses described above, we cultivated PHBV degraders directly from the PHBV trace material of the highest PHBV degrading soil, 2CO (***Fig. 1 – Step 5***, right panel) – using the same PHBV trace material that was scraped from the sensor and used for the cultivation-independent analyses. This effort to cultivate bacteria directly from the degraded PHBV sensors yielded 2 distinct isolates that made PHBV clearing zones *in vitro* (***Fig. 5 inset***). We confirmed their identities through full length 16S rRNA sequencing as *Streptomyces* sp. and *Lysinibacillus* sp. Genomic analyses also identified that both isolates have multiple putative genes associated with PHBV degradation (***Supp. Table 2***). These two taxa were also detected on the PHBV sensors in the cultivation-independent analyses—including in the 2CO soil, where their corresponding ASVs accounted for 0.013% to 0.325% of the bacterial communities—but were too rare to be included in the enrichment analyses described above. This was not unexpected as cultivation-dependent approaches do not necessarily capture the most abundant taxa in any given environment and are often biased towards relatively rare taxa that are amenable to rapid growth under laboratory conditions (Anguita-Maeso *et al*., 2020; Mendes *et al*., 2011). Future work could use a range of different cultivation conditions and media types for the targeted isolation of a broader range of taxa identified from the cultivation-independent analyses described above to test their PHBV degradation potential.

Finally, we tested one of the isolates confirmed to degrade PHBV *in vitro* (*Lysinibacillus* sp.) to corroborate that the degradation was associated with catabolism to CO_2_, not just depolymerization (Lear *et al*., 2021 & 2022). We cultured the *Lysinibacillus* sp. in M9 with the PHBV-PMMA sensor as the sole carbon source, measuring CO_2_ production throughout a five-week incubation (***Fig. 5***). We observed that as the PHBV degradation signal increased over the duration of the experiment, so did cumulative CO₂ production, while PHBV degradation and cumulative CO₂ production were far lower and relatively unchanged over time in the uninoculated controls (***Fig. 5***). These results highlight that the *in situ* PHBV-PMMA sensors reliably capture changes in microbial PHBV degradation (as mentioned above) and that the *Lysinibacillus* sp. isolate is capable of catabolizing PHBV to CO₂.

## Conclusions

We demonstrate a robust strategy to identify putative plastic degrading microbes in soils. We screened 12 distinct soils from across the continental United States using real-time *in situ* sensors and found five with high PHBV degrading capacity. We then used cultivation-independent and cultivation-dependent approaches for the targeted identification of novel PHBV-degrading microbes. From DNA extracted directly from the PHBV trace material of sensors incubated in high PHBV degrading soils, we used marker gene sequencing to identify putative bacterial and fungal taxa enriched on PHBV. We also cultured novel PHBV degrading isolates of *Lysinibacillus* sp. and *Streptomyces* sp. directly from the PHBV trace material and used the sensors to demonstrate their plastic degrading capacity *in vitro*. From our targeted approach, we found i) PHBV degradation rates vary across soils in ways that are difficult to predict from *a priori* information on soil characteristics, ii) pairing real-time measurements of PHBV degradation with analyses of the microbes colonizing the PHBV makes it possible to identify putative PHBV degrading microbes in soil, including some taxa confirmed to be capable of PHBV degradation via *in vitro* assays, and iii) the diversity of soil microbes capable of degrading PHBV is likely higher than expected from isolate screening efforts, as many of these taxa have not previously been identified as plastic degraders from more traditional cultivation-based screening efforts.

Going forward, our strategy could be expanded to target a broader range of biodegradable and emerging polymers by modifying the plastic substrate used in the degradable trace. This would enable the development of *in situ* sensors that not only identify microbial hotspots of degradation, but also directly assess the environmental biodegradability of different plastics *in situ*. Deploying these sensors across diverse environments, including soils, composting systems, and landfills, would provide a powerful platform for discovering novel plastic degraders and evaluating the fate of plastic materials outside the laboratory. Furthermore, coupling real-time, continuous sensor-based measurements with manipulative experiments could reveal how environmental factors such as temperature, moisture, and nutrient availability, influence degradation rates, and the microbial taxa responsible. Ultimately, expanding our new targeted strategy will help unlock Earth’s vast, unexplored capacity for plastic degradation, improve our ability to predict and enhance plastic degradation in natural systems, and provide new tools to mitigate the escalating plastic pollution crisis.

## Acknowledgements

AJCB thanks Cliff Bueno de Mesquita and Elias Stallard-Olivera for help collecting soils, Caihong Vanderburgh for guidance on ONT sequencing, and the Fond de Recherche du Québec – Nature et Technologies for salary support (Bourses de recherche postdoctorale 346908). Funding for this work was provided by grants to NF and GW from the US Department of Defense Army Research Office (W911NF-26-1-A013) and the Research and Innovation Office of the University of Colorado Boulder.

---

**Supp.Table 1.** Soil information and selected edaphic variables.

|  | 1CO | 2CO | 3CO | 1MA | 2MA | 1MI | 2MI | 1CA | 2CA | 3CA | 4CA | 5CA |
| --- | --- | --- | --- | --- | --- | --- | --- | --- | --- | --- | --- | --- |
| Soil Descriptor | Montane | Community Garden | Subalpine | Forest | Campus Field | Conventional Farm | Regenerative Farm | Campus Experimental Smart Farm | Fallow Urban Farm | Regenerative Urban Farm | Elementary School Lawn | Grassland Reserve |
| Location | Betasso, CO | Boulder, CO | Nederland, CO | Mount Holyoke, MA | Mount Holyoke, MA | Kellogg Biological Station, MI | Kellogg Biological Station, MI | Merced, CA | Fremont, CA | Fremont, CA | Oakland, CA | Merced, CA |
| Latitude | 40°01'05.8"N | 40°00'05.7"N | 39°56'48.2"N | 42°15'7"N | 42°15'15"N | 42°24'17"N | 42°24'17"N | 37°21'19"N | 37°34'19"N | 37°34'20"N | 37°48'26"N | 37°22'30"N |
| Longitude | 105°20'40.5"W | 105°13'58.8"W | 105°30'49.9"W | 72°34'17"W | 72°34'37"W | 85°22'44"W | 85°22'47"W | 120°24'51"W | 121°58'09"W | 121°58'10"W | 122°17'51"W | 120°24'33"W |
| Elevation (m) | 2004.06 | 1655.064 | 2661.208 | 90.1 | 65.0 | 280.1 | 277.0 | 65.3 | 23.9 | 23.0 | 6.0 | 86.2 |
| Collection Date | March, 2025 | March, 2025 | March, 2025 | July, 2025 | July, 2025 | May, 2024 | May, 2024 | July, 2025 | July, 2025 | July, 2025 | July, 2025 | July, 2025 |
| Mean Respiration<br>($\mu\text{g C-CO}_2\text{ g Soil}^{-1}\text{ Day}^{-1}$) | 5.442113 | 11.984232 | 4.921061 | 4.854334 | 6.681851 | 10.508209 | 4.054799 | 16.622768 | 15.631607 | 11.405282 | 15.573713 | 26.618888 |
| pH | 6.7 | 7.4 | 6.5 | 5.1 | 6.6 | 5.9 | 5.7 | 6.3 | 7.2 | 7.0 | 5.8 | 6.0 |
| Soluble Salts<br>(mmho $\text{cm}^{-1}$) | 0.17 | 0.75 | 0.21 | 0.10 | 0.26 | 0.29 | 0.40 | 0.80 | 0.43 | 0.69 | 0.51 | 0.27 |
| % Organic Matter | 4.9 | 13.9 | 1.8 | 6.4 | 5.6 | 1.2 | 1.6 | 6.8 | 10.4 | 14.1 | 8.0 | 5.5 |
| NO<sub>3</sub>-N (ppm) | 7.7 | 71.4 | 11.2 | 9.4 | 23.9 | 26.7 | 40.6 | 81.6 | 21.9 | 63.4 | 27.7 | 4.0 |
| P (ppm) | 24 | 702 | 36 | 179 | 47 | 99 | 49 | 37 | 136 | 100 | 71 | 16 |
| K (ppm) | 200 | 560 | 88 | 45 | 72 | 128 | 144 | 266 | 175 | 235 | 299 | 211 |
| Ca (ppm) | 1455 | 2888 | 875 | 56 | 1237 | 358 | 503 | 2219 | 3283 | 2355 | 1862 | 1282 |
| Mg (ppm) | 175 | 700 | 102 | 21 | 66 | 49 | 79 | 619 | 477 | 449 | 296 | 357 |
| Na (ppm) | 8 | 45 | 4 | 5 | 8 | 2 | 3 | 52 | 31 | 65 | 41 | 24 |
| Cation Exchange<br>Capacity<br>(me 100 g<sup>-1</sup>) | 9.3 | 21.9 | 5.7 | 7.0 | 7.0 | 4.0 | 4.9 | 17.3 | 21.0 | 16.4 | 12.7 | 12.9 |
| SO<sub>4</sub> (ppm) | 8.0 | 50.1 | 3.3 | 18.9 | 14.2 | 10.5 | 9.2 | 9.2 | 20.0 | 18.0 | 17.0 | 14.7 |

**Supp.Table 2.** Genomic Analysis of PHBV Degrading Isolates for Known Degradation Genes.

| Plastic DB<br>Protein ID | Streptomyces sp.<br>[Garden]<sup>a</sup> | Lysinibacillus sp.<br>[Garden]<sup>a</sup> | Degradation Gene \|\| Reference Organism \|\| Polymers<sup>b</sup> |
| --- | --- | --- | --- |
| 00211 | 2 | 1 | Lipase \|\| *Pseudomonas chlororaphis* \|\| PHBV, PHA, PLA, PCL, PES |
| 00074 | 1 | 0 | Cutinase \|\| *Saccharomonospora viridis* \|\| PBAT, PBS, PBSA, PCL, PET, PHB, PLA, PHA |
| 00071 | 4 | 4 | 3HV_dehydrogenase \|\| *Paracoccus denitrificans* \|\| P3HV, PHBV, PHA |
| 00007 | 1 | 0 | PHB_depolymerase \|\| *Paucimonas lemoignei* \|\| PHB, PHA |
| 00048 | 0 | 1 | PLA_depolymerase \|\| Uncultured bacterium \|\| PBS, PBSA, PCL, PES, PLA, PHA |
| 00072 | 0 | 1 | PHB_depolymerase \|\| *Bacillus thuringiensis* \|\| PHB, PHA |

<sup>a</sup>, Number of hits in the genome of the isolate to the identified degradation gene
<sup>b</sup>, PLA, polylactic acid; PCL, polycaprolactone; PES, polyethersulfone; PBAT, polybutylene adipate terephthalate; PBS, polybutylene succinate;
PBSA, polybutylene succinate-co-adipate; P3HV, poly(3-hydroxyvalerate).

**Supp. Fig. 1.**
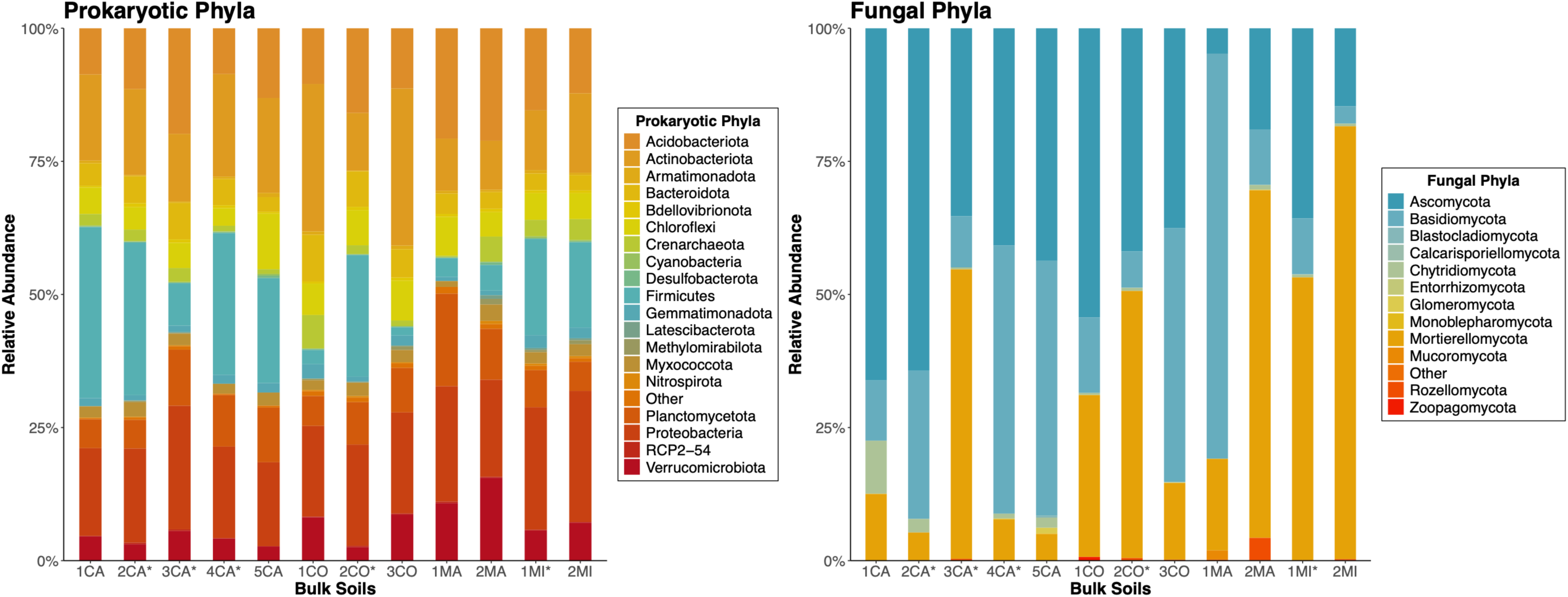
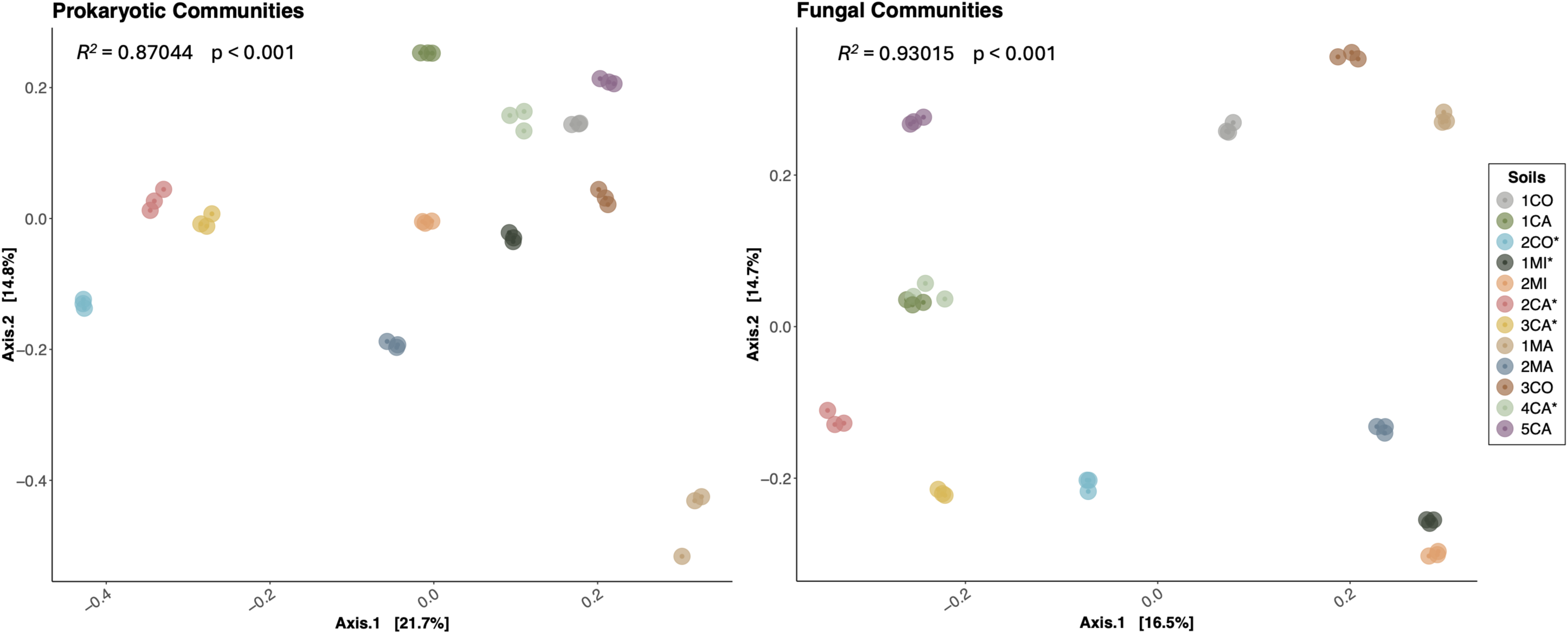
16S rRNA gene and ITS sequencing results showing that each of the 12 soils screened for PHBV degradation have distinct microbial communities. A) Relative abundance of prokaryotic (left) and fungal (right) phyla among the 12 soils. Prokaryotic phyla that were represented below 0.03 % reads, or 0.0005% for fungi, were regrouped as “Other”. B) Principal component analysis of Bray-Curtis distances between the 12 soils captured 36.5%, and 31.2%, of the variance among the prokaryotic (left), and fungal (right), communities, respectively. PERMANOVA identified soil types as a significant experimental factor (Prokaryotes, R^2^ = 0.87044, p < 0.001; Fungi, R^2^ = 0.93015, p < 0.001). Soils used for subsequent microbial analysis of potential PHBV degraders are marked with asterisks.

**Supp Fig. 2.**
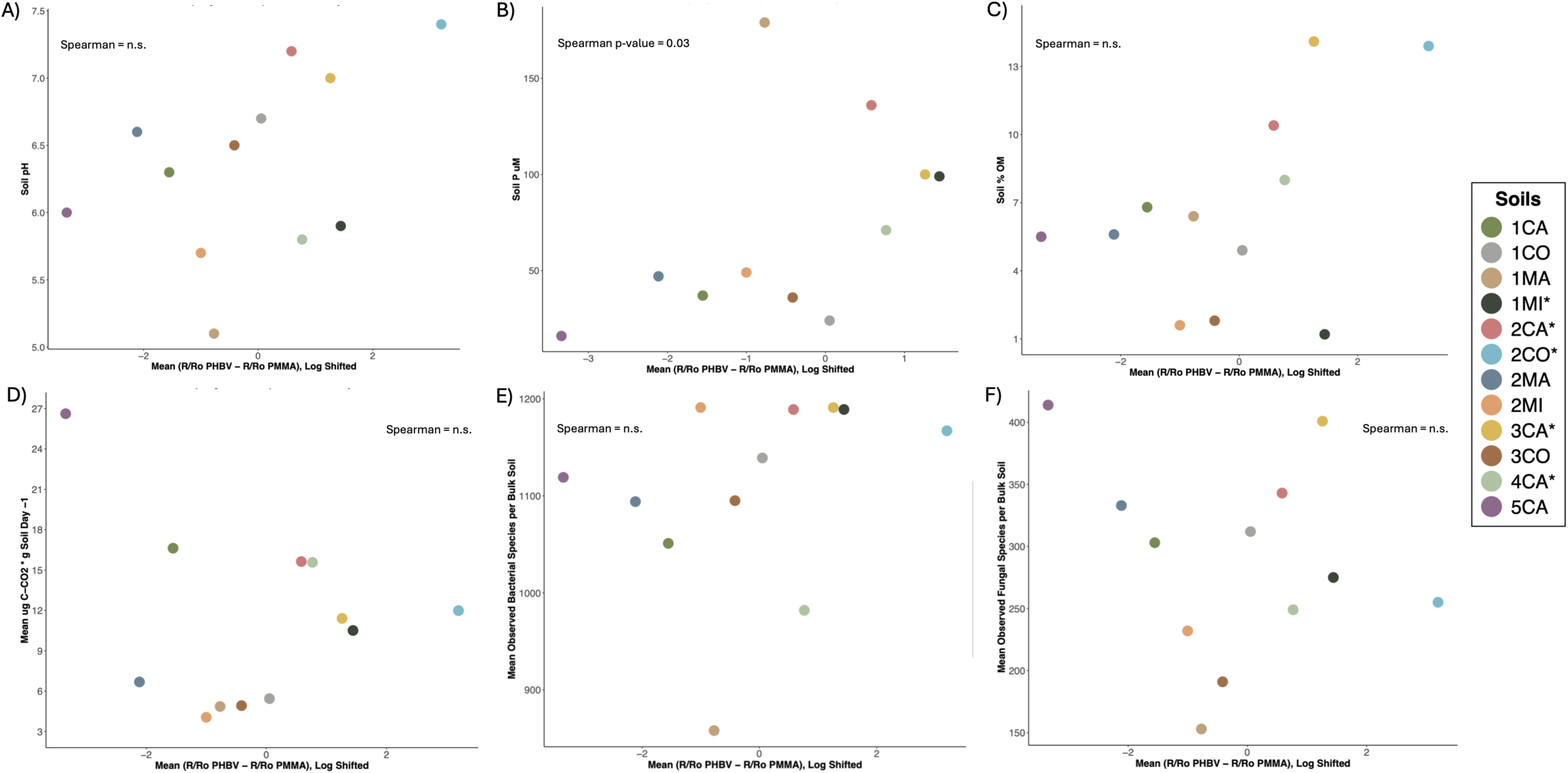
PHBV degradation was not strongly correlated with other. A-C) soil edaphic characteristics (pH, P, %OM), D) microbial activity (measured as µg C-CO_2_ g Soil^-1^ Day^-1^), E) mean bacterial species richness (858 – 1291), nor F) mean fungal species richness (153 – 401). We used a Spearman rank correlation. The only significant (p = 0.03, rho = 0.64) correlation identified was in B) P vs degradation signal, due to the high P concentration in the 2CO sample. Soils were rarified by type to determine bacterial and fungal species richness.

**Supp. Fig. 3.**
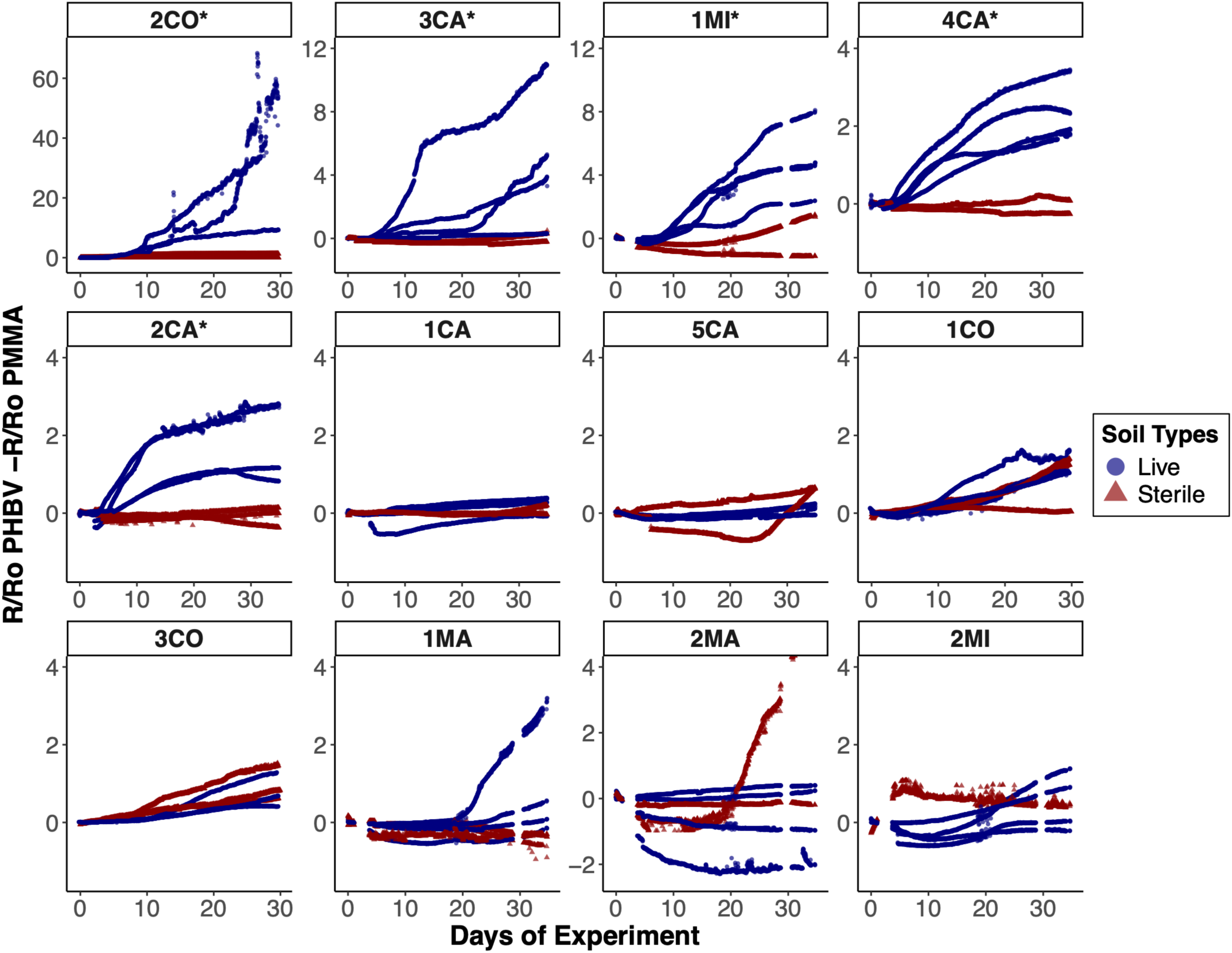
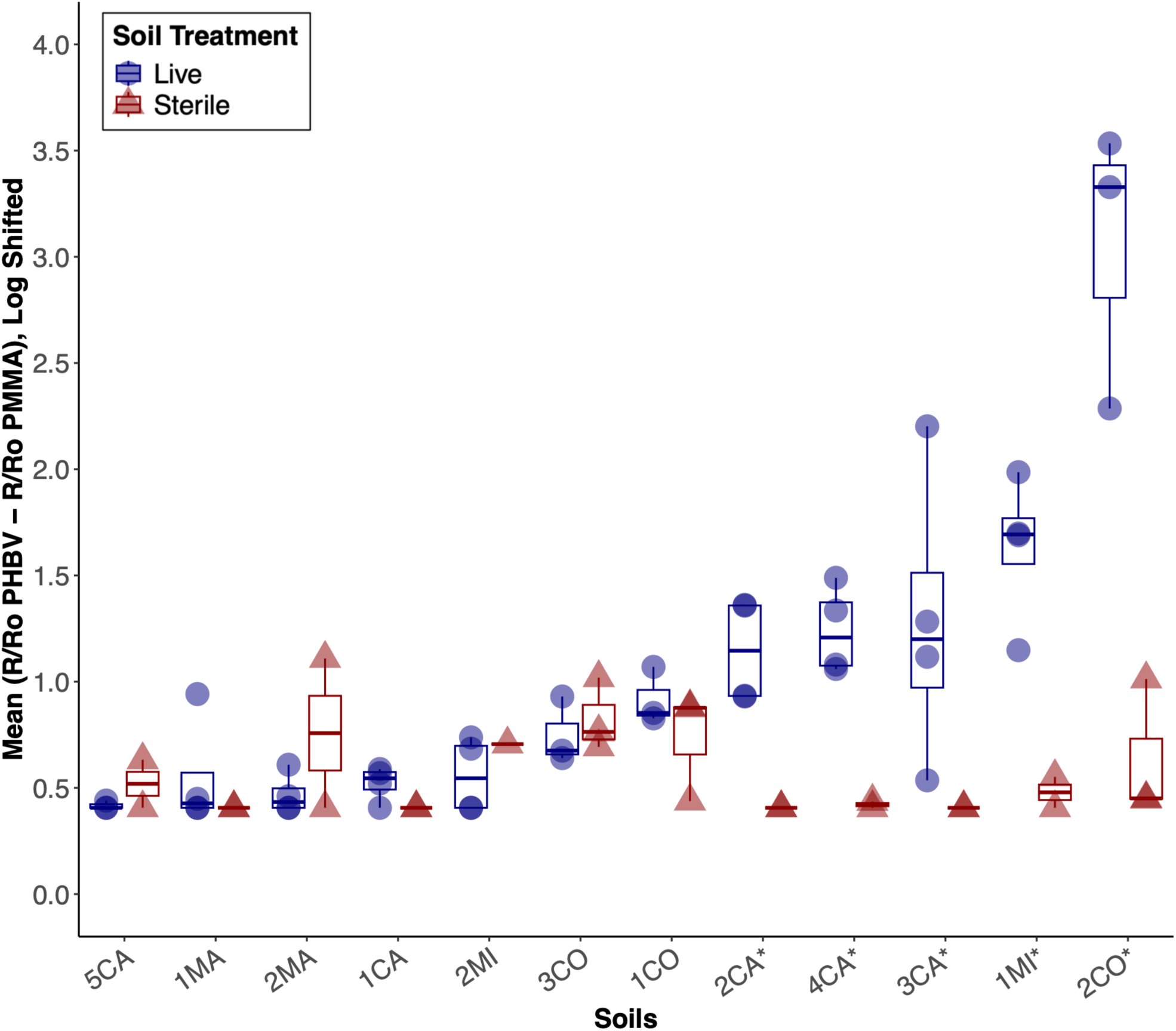
PHBV degradation signals from the sterilized subsamples of the 12 screened soils collected from across the USA (CA, California; CO, Colorado; MA, Massachusetts; MI, Michigan). Soil mesocosms were prepared as live soils, or triple autoclaved sterile soils. A) Change in PHBV degradation signals over time across the 5-week incubation period in live (blue) and sterile (red) soils. B) Day 19.5-29.5 summarized PHBV degradation sensor signal for live (blue circles) and sterilized (red triangles) soil mesocosms, as the highest signal was recorded from this timeframe due to accumulated degradation. N = 4 for all live soil mesocosms presented here, except 1CO, 2CO, 3CO, and 5CA, which had 3. We removed one 5CA sensor from the data due to mechanical damage. N = 2 for all sterile soil mesocosms, except 1CO, 2CO, 3CO, and 5CA, which had 3. We removed one 2MI sterile sensor from the dataset due to known contamination with live soil.

**Supp. Fig. 4.**
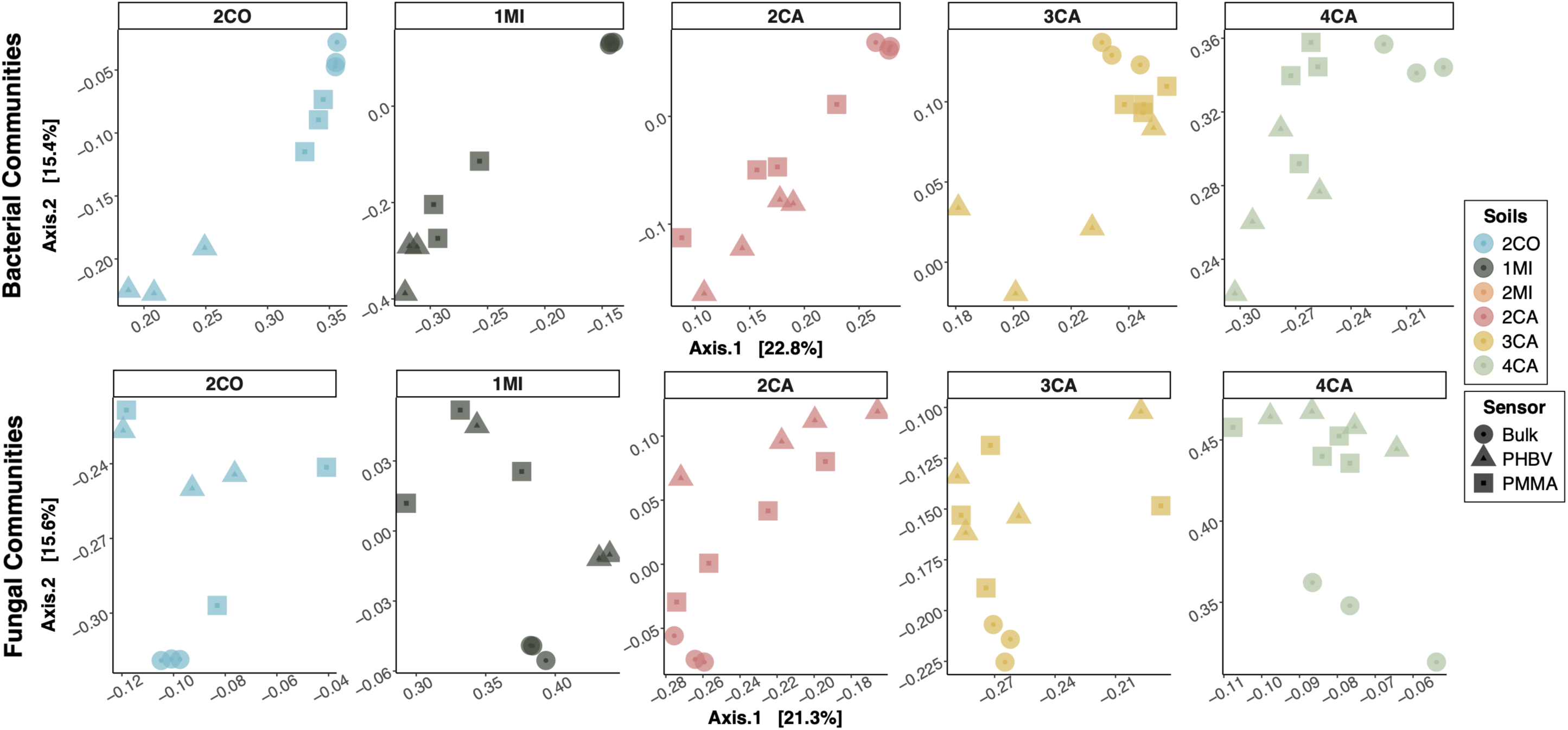
Prokaryotic (top row) and fungal (bottom row) communities in the five strongest PHBV degrading soils are significantly different (PermANOVA p < 0.01) between bulk soils and their corresponding PHBV and PMMA communities. However, the PHBV and PMMA communities were not consistently significantly different from each other. We rarefied each of the five soil types independently, and calculated a Bray-Curtis dissimilarity matrix for prokaryotes and fungi for each soil, before replotting them as PCoAs.

